# *fruitless* tunes functional flexibility of courtship circuitry during development

**DOI:** 10.1101/2020.06.20.163055

**Authors:** Jie Chen, Sihui Jin, Jie Cao, Qionglin Peng, Yufeng Pan

## Abstract

*Drosophila* male courtship is controlled by the male-specific products of the *fruitless (fru^M^* gene and its expressing neuronal circuitry. *fru^M^* is considered a master gene that controls all aspects of male courtship. By temporally and spatially manipulating *fru^M^* expression, we found that *fru^M^* is required during a critical developmental period for innate courtship towards females, and its function during adulthood is relatively trivial. By altering or eliminating *fru^M^* expression, we generated males that are innately heterosexual, homosexual, bisexual, or without innate courtship but could acquire such behavior in an experience-dependent manner. These findings show that *fru^M^* is not absolutely necessary for courtship but is critical during development to build a sex circuitry with reduced flexibility and enhanced efficiency and provide a new view about how *fru^M^* tunes functional flexibility of a sex circuitry.

## Introduction

*Drosophila* male courtship is one of the best understood innate behaviors in terms of genetic and neuronal mechanisms (Dickson, 2008; Yamamoto and Koganezawa, 2013). It has been well established that the *fruitless (fru)* gene and its expressing neurons control most aspects of such innate behavior (Ito et al., 1996; Manoli et al., 2005; Ryner et al., 1996; Stockinger et al., 2005). The male-specific products of the P1 promoter of the *fru* gene *(fru^M^)* are expressed in ~2000 neurons, which are interconnected to form a sex circuitry from sensory neurons to motor neurons (Cachero et al., 2010; Lee et al., 2000; Manoli et al., 2005; Stockinger et al., 2005; Usui-Aoki et al., 2000; Yu et al., 2010). *fru^M^* function is necessary for the innate courtship behavior and sufficient for at least some aspects of courtship (Baker et al., 2001; Demir and Dickson, 2005; Manoli et al., 2005). Thus, the study of *fru^M^* function in controlling male courtship serves as an ideal model to understand how innate complex behaviors are built into the nervous system by regulatory genes (Baker et al., 2001).

Although *fru^M^* serves as a master gene controlling *Drosophila* male courtship, we recently found that males without *fru^M^* function, although do not court if raised in isolation, were able to acquire at least some courtship behaviors if raised in groups (Pan and Baker, 2014). Such *fru^M^*-independent but experience-dependent courtship acquisition requires another gene in the sex determination pathway, the *doublesex (dsx)* gene (Pan and Baker, 2014). *dsx* encodes male- and female-specific DSX proteins (DSX^M^ and DSX^F^, respectively) (Burtis and Baker, 1989), and DSX^M^ is expressed in *~*700 CNS neurons, the majority of which also express *fru^M^* (Rideout et al., 2010; Robinett et al., 2010). It has been found that the *fru^M^* and *dsx^M^* coexpressing neurons are required for courtship in the absence of *fru^M^* function (Pan and Baker, 2014). Thus *fru^M^*-expressing neurons, especially those co-expressing *dsx^M^*, control the expression of courtship behaviors even in the absence of FRU^M^ function. Indeed, although the gross neuroanatomical features of the *fru^M^*-expressing circuitry are largely unaffected by the loss of *fru^M^,* detailed analysis revealed morphological changes of many *fru^M^*-expressing neurons (Kimura et al., 2005; Kimura et al., 2008; Mellert et al., 2010). Recent studies further reveal that FRU^M^ specifies neuronal development by recruiting chromatin factors and changing chromatin states, and also by turning on and off the activity of the transcription repressor complex (Ito et al., 2012; Ito et al., 2016; Sato et al., 2019a; Sato et al., 2019b; Sato and Yamamoto, 2020).

That FRU^M^ functions as a transcription factor to specify development and/or physiological roles of certain *fru^M^*-expressing neurons, and perhaps the interconnection of different *fru^M^*-expressing neurons to form a sex circuitry raises important questions regarding when *fru^M^* functions and how it contributes to the sex circuity *(e.g.,* how the sex circuitry functions differently with different levels of FRU^M^), especially in the background that *fru^M^* is not absolutely necessary for male courtship (Pan and Baker, 2014). To at least partially answer these questions, we temporally or spatially knocked down *fru^M^* expression, and compared courtship behavior in these males with that in wild-type males or *fru^M^* null males and revealed crucial roles of *fru^M^* during a narrow developmental window for the innate courtship towards females. We also found that the sex circuitry with different *fru^M^* expression has distinct function such that males could be innately heterosexual, homosexual, bisexual, or without innate courtship but could acquire such behavior in an experience-dependent manner. Thus, *fru^M^* tunes functional flexibility of the sex circuitry instead of switching on its function as conventionally viewed.

## Results

To specifically knock down *fru^M^* expression, we used a microRNA targeting *fru^M^ (UAS-fruMi)* and a scrambled version as a control *(UAS-fruMiScr)* as previously used (Chen et al., 2017). We firstly tested male courtship without food in the behavioral chamber. Knocking down *fru^M^* in all the *fru^GAL4^* labeled neurons eliminated male courtship towards females (courtship index [CI], which is the percentage of observational time that males displayed courtship, is nearly 0) (Figure 1A), consistent with previous findings that *fru^M^* is required for innate male-female courtship (Demir and Dickson, 2005; Pan and Baker, 2014). We then added a temperature dependent *tub-GAL80^ts^* transgene to restrict *UAS-fruMi* expression *(e.g.,* at 30°C) at different developmental stages. We raised *tub-GAL80^ts^/+; fru^GAL4^/UAS-fruMi* flies at 18°C (permissive for GAL80^ts^ that inhibits GAL4 activity) and transferred these flies to fresh food vials every two days. In this way we generated *tub-GAL80 ^ts^/+; fru^GAL4^/UAS-fruMi* flies at 9 different stages from embryos to adults and incubated all flies at 30°C to allow *fru^M^* knock-down for 2 days, then placed all flies back to 18°C until courtship test (Figure 1B). We found that males with *fru^M^* knocked down from stage 5 to 6, matching the pupation phase, rarely courted (CI < 10%) and none successfully mated, while males with *fru^M^* knocked down near this period showed a partial courtship deficit (Figure 1C, D). Knocking down *fru^M^* specifically during adulthood for 2 days did not affect male courtship (CI > 80%) and mating success. These results reveal a critical developmental period where *fru^M^* is required for adult male courtship towards females.

**Figure 1:**
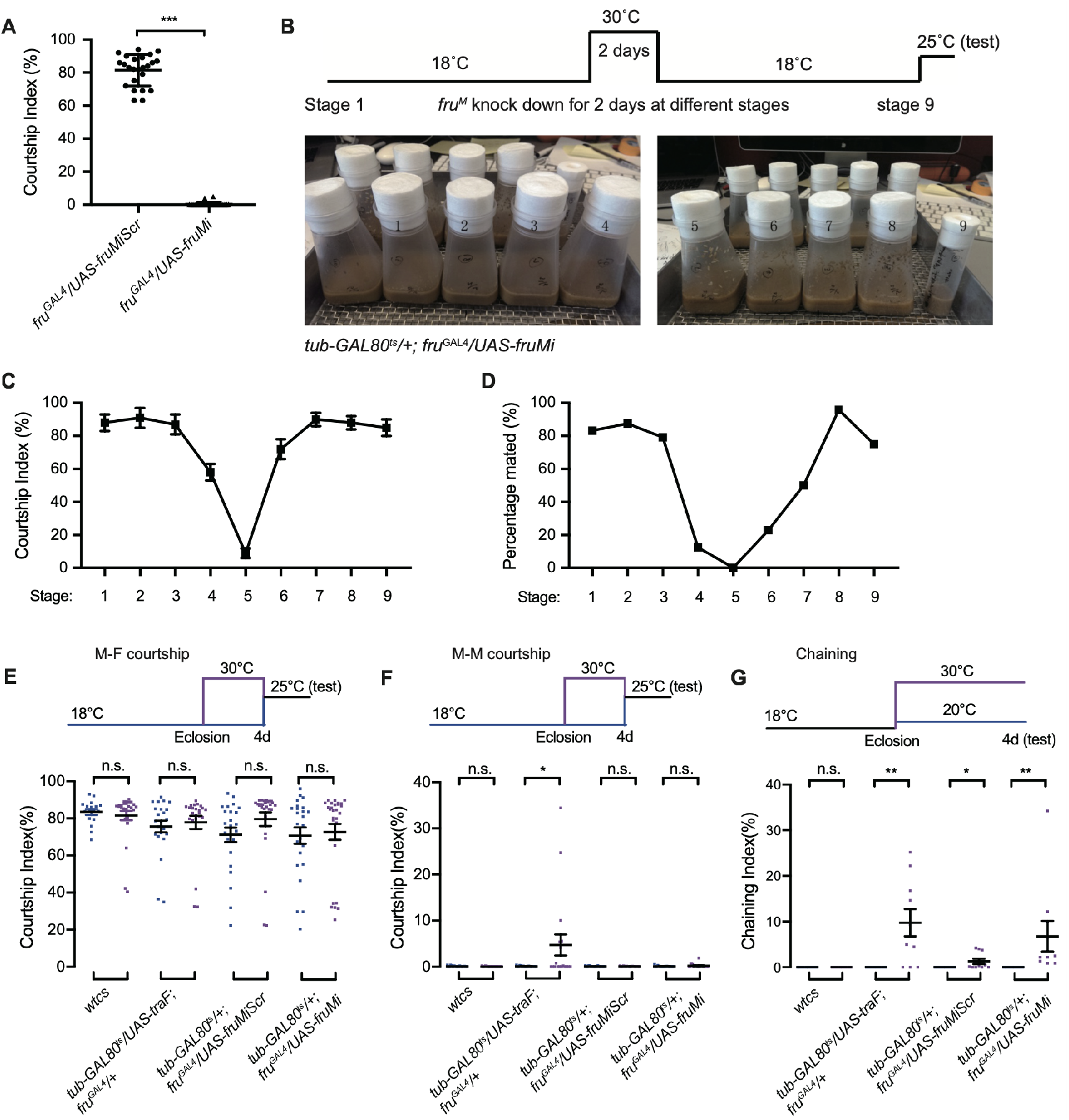
Distinct roles of *fru^M^* during development and adulthood in regulating male courtship. **A:** Knocking down *fru^M^* using RNAi throughout development and adulthood eliminated male courtship towards virgin females. *n* = 24 for each. \*\*\**p* < 0.001, unpaired t-test. **B**: A schematic of genetic strategy to knock down *fru^M^* at different developmental stages for 2 days. Stages 1 to 9 refer to specific developmental stages from embryos to adults with interval of 2 days. **C** and **D**: Courtship indices of males with *fru^M^* knocked down at specific developmental stages as indicated above towards virgin females. Males with *fru^M^* knocked down at stage 5 for 2 days (a period of pupation from stage 5 to 6, see above picture) rarely courted virgin females (**C**), and none successfully mated (**D**). Knocking down *fru^M^* at stages near 5 (*e.g.*, stage 4 or 6) also partially impairs courtship and mating success. Knocking down *fru^M^* at stage 9 (adult) has no obvious effect on courtship and mating. *n* = 24 for each. **E-G**: Knocking down *fru^M^* specifically during adulthood slightly increased male-male courtship behaviors. For male-female courtship (**E**), *n* = 17, 26, 23, 23, 24, 27, 24 and 28 respectively (from left to right), n.s., not significant, unpaired t-test. For single-pair male-male courtship (**F**), *n* = 18 for each. n.s., not significant, \**p* < 0.05, unpaired t-test. For male chaining among 8 males as a group (**G**), *n* = 8, 8, 8, 10, 8, 9, 8 and 9 respectively (from left to right). n.s., not significant, \**p* < 0.05, \*\**p* < 0.01, Mann-Whitney test. Error bars indicate SEM.

As we did not see an obvious courtship deficit in males with *fru^M^* knocked down in adulthood for 2 days, we further tested the role of *fru^M^* in adulthood using different approaches. We set out to express the female-specific *transformer (traF)* gene (Baker and Ridge, 1980; McKeown et al., 1988) to feminize all *fru^GAL4^* labeled neurons, in addition to the *fru^M^* RNAi experiments. We express *UAS-traF* or *UAS-fruMi* in all the *fru^GAL4^* labeled neurons specifically during adulthood for 4 days before test (see procedure above each figure) for single-pair male-female, male-male, and male chaining (in groups of 8 males) behaviors. We found that overexpression of *traF* in all *fru^GAL4^* labeled neurons during adulthood for 4 days did not affect male-female courtship (Figure 1E), but slightly increased male-male (Figure 1F) and male chaining behaviors (Figure 1G). Furthermore, knocking down *fru^M^* in all *fru^GAL4^* labeled neurons during adulthood for 4 days did not affect male-female (Figure 1E) or male-male courtship (Figure 1F), but slightly increased male chaining behaviors (Figure 1G). Together these results indicate that *fru^M^* function during pupation is crucial for adult courtship towards females, while its function during adulthood is dispensable for female-directed courtship, though it plays a minor role in inhibiting male-male courtship behaviors.

To further reveal the role of *fru^M^* in male courtship, we tried to spatially knock down *fru^M^* expression using a simple way: *fru^M^* in brain and *fru^M^* outside brain. We used *Otd-Flp* expressing FLP specifically in the central brain (Asahina et al., 2014) to divide *fru^GAL4^* expression (Figure 2A) into two parts: *fru^M^*- and *Otd*-positive neurons (specifically in brain) in *Otd-Flp/UAS-myrGFP;fru^GAL4^/tub>GAL80>* males (Figure 2B), and *fru^M^*-positive but *Otd*-negative neurons (theoretically outside brain, but still with few in brain) in *Otd-Flp/UAS-myrGFP; fru^GAL4^/tub>stop>GAL80* males (Figure 2C). We also checked GFP expression in peripheral nervous system in these males, and found a few GFP-positive cells in antennae and forelegs in *Otd-Flp/UAS-myrGFP; fru^GAL4^/tub>stop>GAL80* males, but no expression in *Otd-Flp/UAS-myrGFP; fru^GAL4^/tub>GAL80>* or wild-type males (Figure 2D-G). Thus, we successfully divided *fru^GAL4^* expression into two categories, one with *GAL4* expressed in *fru^+^Otd^+^* neurons in brain, and the other with *GAL4* expressed in *fru^+^Otd^-^* neurons outside brain.

**Figure 2:**
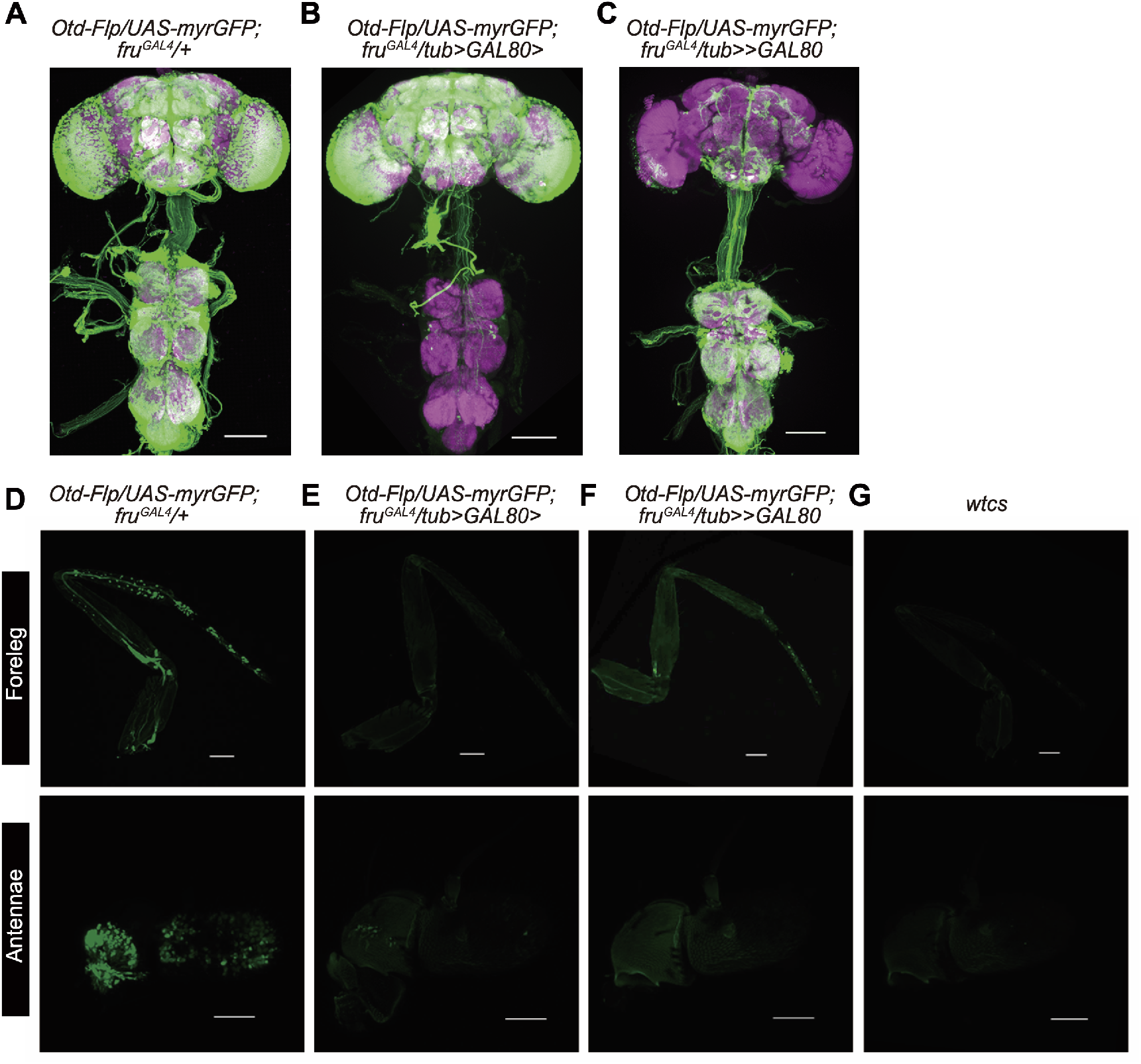
Dividing *fru^M^* expression into two complementary parts. **A**: Expression pattern of the *fru^M^* circuitry revealed by *fru^GAL4^* driving *UAS-myrGFP* (green), costained with nc82 (magenta). **B** and **C**: Genetic strategy to divide *fru^M^* neurons (green) into two parts. Scale bars above, 100μm. **D-G**: Expression pattern of *fru^GAL4^* driving *UAS-myrGFP* (green) in forelegs (scale bars, 100 mm) and antennae (scale bars, 50 mm) in males with indicated genotypes. Representative of 5 samples each.

We then used the above intersectional strategy to knock down *fru^M^* expression in all *fru^GAL4^* neurons or those neurons in or outside brain, and compared male courtship with that in wild-type males and *fru^M^* null males. We tested one-time single-pair male-female and male-male courtship (single-housed before test) as well as male chaining in groups of 8 males over 3 days on food for better comparison of these courtship assays, as courtship by *fru^M^* null males largely depends on food presence (Pan and Baker, 2014). We found that male-male courtship in *fru^M^* knocked down males is higher if tested on food, consistent with a courtship promoting role by food (Grosjean et al., 2011; Pan and Baker, 2014), while courtship in wild-type males on food or without food is not changed in our assays (Figure 3—figure supplement 1).

We found that wild-type males performed intensive courtship behavior towards virgin females (CI > 80%) and rarely courted males (CI ~0) (Figure 3A). Furthermore, these control males did not show any chaining behavior after grouping from 3 hours to 3 days (ChI = 0) (Figure 3B). In striking contrast, *fru^M^* null mutant *(fru^LexA^/fru^4-40^)* males rarely courted either females or males (Figure 3C); however, these males developed intensive chaining behavior after grouping for 1-3 days (Figure 3D). These observations replicated previous findings that there exists a *fru^M^*-independent experience- and *dsx^M^*-dependent courtship pathway (Pan and Baker, 2014) (Figure 3E). To compare behavioral differences by *fru^M^* null males and *fru^M^* RNAi knocked down males, we quantified to how much extent the microRNA against *fru^M^* worked. We found that the *fru^M^* mRNA level was reduced to ~40% of that in control males (Figure 3F). Interestingly, while males with *fru^M^* knocked down in all *fru^M^* neurons rarely courted females (CI~5%, Figure 3G), they displayed a high level of male-male courtship behavior (CI > 50%, Figure 3G) and constantly high level of male chaining (Figure 3H), dramatically different from *fru^M^* null males. These results reveal distinct roles of low *fru^M^* (RNAi) and high *fru^M^* (wild-type) in regulating male-male and male-female courtship (Figure 3I). To further dissect the role of *fru^M^* in male courtship, we knocked down *fru^M^* specifically in brain, and found that such males had a reduced level of courtship towards females (CI = 56.61 ± 5.86%), but their sexual orientation was not changed as they courted males in a much lower level (CI = 15.94 ± 3.26%, Figure 3J). Furthermore, males with *fru^M^* knocked down in brain showed low male chaining behavior initially but increasing levels of chaining behavior over 1-3 days (ChI[3h] = 9.35 ± 5.40%, ChI[3d] = 68.82 ± 5.53%, Figure 3K). These results indicate that *fru^M^* function in brain promotes male-female courtship and inhibits acquisition or progression of the experience-dependent chaining behavior (Figure 3L). In contrast, males with *fru^M^* knocked down outside brain showed equally intensive male-female and male-male courtship (CI[male-female] = 85.62 ± 1.42%, CI[male-male] = 82.89 ± 2.76%, Figure 3M), indicating an inhibitory role of *fru^M^* in these neurons for male-male courtship (Figure 3O). These males performed a high level of male chaining behavior initially (ChI[3h] =92.90 ± 3.08%), but decreased levels of chaining behavior over 1-3 days (ChI[3d] = 20.01 ± 3.75%, Figure 3N), consistent with the above finding that *fru^M^* function in the brain which is intact in these males inhibits acquisition or progression of male chaining behavior (Figure 3L).

**Figure 3:**
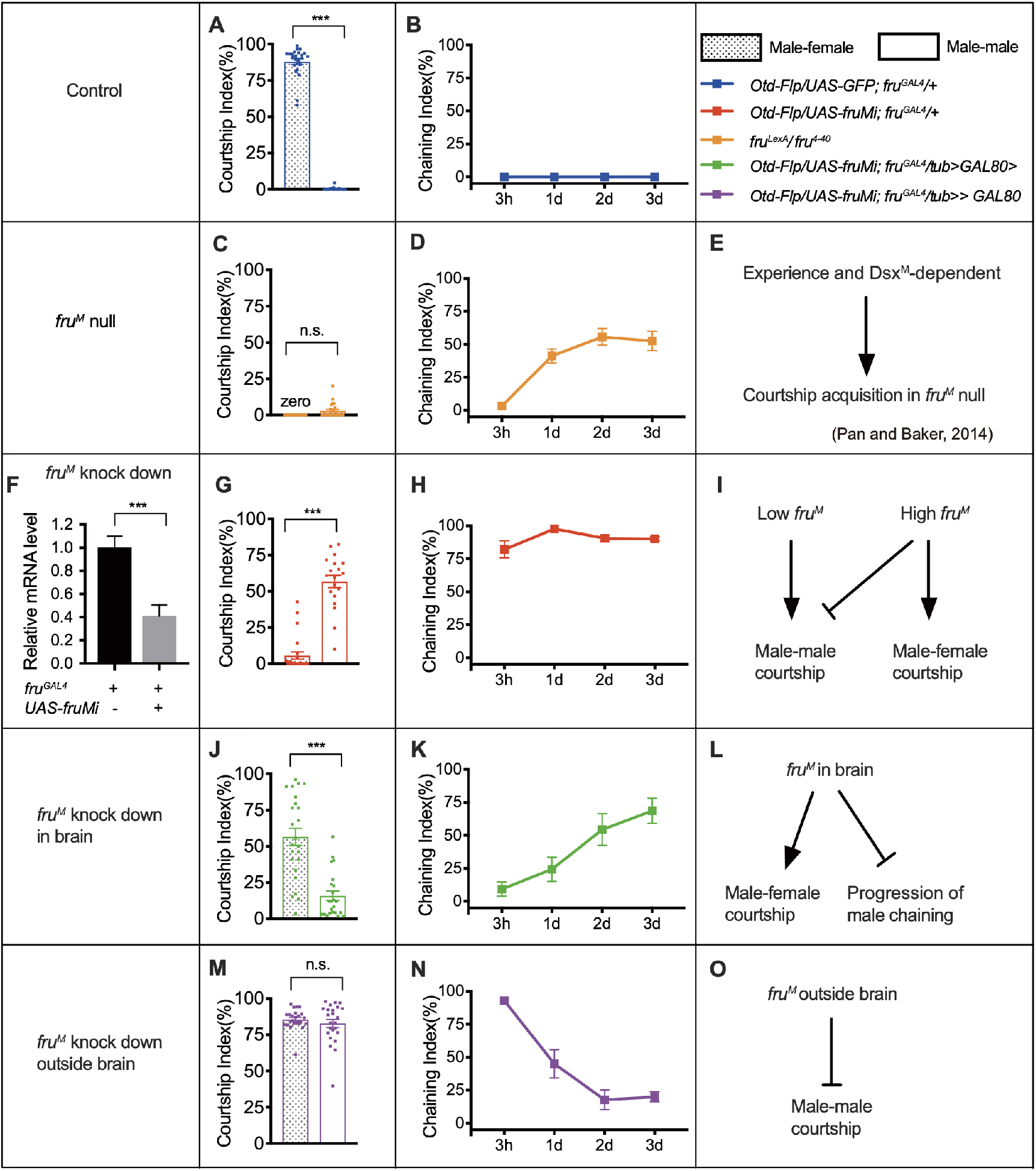
*fru^M^* tunes functional flexibility of the *fru^M^* circuitry. **A** and **B**: Wild-type males courted intensively towards virgin females (**A**, left bar), but rarely courted males (**A**, right bar) or displayed chaining behavior in groups of 8 males (**B**). *n* = 24, 24, 8 respectively. \*\*\**p* < 0.001, unpaired t-test. **C**: *fru^LexA^/fru^4-40^* (*fru^M^* null) males rarely courted either females or males. *n* = 24 for each. n.s., not significant, unpaired t-test. **D**: *fru^LexA^/fru^4-40^* males did not show chaining behavior after 3-hr group-housing, but developed intensive chaining behavior after1-3 days. *n* = 8. **E**: A summary of courtship acquisition independent of *fru^M^*. **F**: RNAi against *fru^M^* efficiently decreased but not fully eliminated *fru^M^* expression. *n* = 3. \*\*\**p* < 0.001, Mann-Whitney U test. **G**: Knocking down *fru^M^* in all *fru^GAL4^* neurons generated males that have reversed sexual orientation such that they rarely courted females but intensively courted males. *n* = 24 and 19 respectively. \*\*\**p* < 0.001, unpaired t-test. **H**: Males with *fru^M^* knocked down in all *fru^GAL4^* neurons showed intensive chaining behavior at all time points (from 3 hours to 3 days upon group-housing). *n* = 7. **I**: Distinct roles of low *fru^M^* (RNAi) and high *fru^M^* (wild-type) in regulating male-male and male-female courtship. **J**: Males with *fru^M^* knocked down in *fru^GAL4^* neurons in the brain had a lower level of courtship towards females, but their sexual orientation was not changed. *n* = 24 and 23 respectively. \*\*\**p* < 0.001, unpaired t-test. **K**: Males with *fru^M^* knocked down in *fru^GAL4^* neurons in brain showed low male chaining behavior initially but increasing levels of chaining behavior over 1-3 days. *n* = 6. **L**: A summary of the role of *fru^M^* in brain in promoting male-female courtship and suppressing the experience-dependent acquisition or progression of male chaining behavior. **M**: Males with *fru^M^* knocked down in *fru^GAL4^* neurons outside brain generated bisexual males that have intensive male-female and male-male courtship. *n* = 24 for each. n.s., not significant, unpaired t-test. **N**: Males with *fru^M^* knocked down in *fru^GAL4^* neurons outside brain showed high male chaining behavior initially, but decreased levels of chaining behavior over 1-3 days. *n* = 8. **O**: A summary of the role of *fru^M^* outside brain in suppressing male-male courtship behavior. Error bars indicate SEM.

Taken together, the above results indicate crucial role of *fru^M^* expression level during a critical developmental period for the manifestation of courtship behaviors and reveal functional flexibility of the *fru^M^*-expressing sex circuitry (Figure 4).

**Figure 4.**
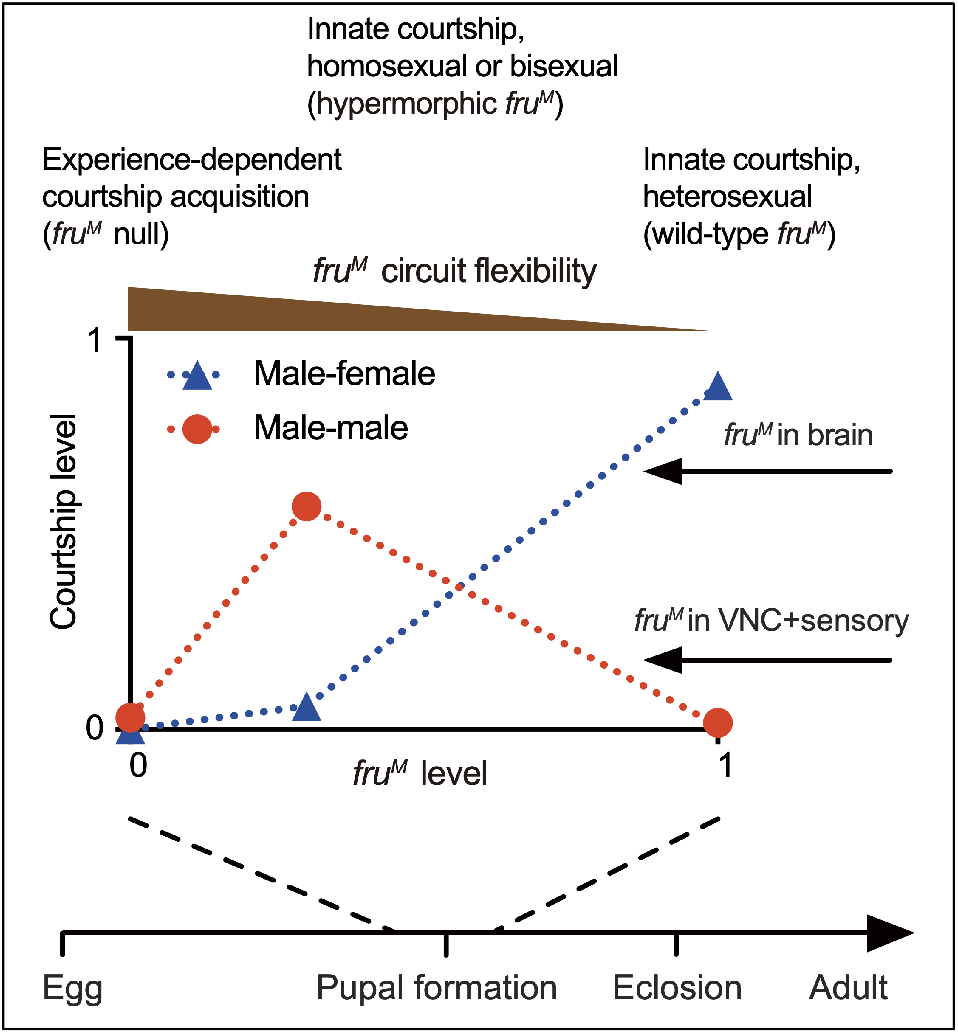
A summary of *fru^M^* function in male courtship. Firstly, *fru^M^* is largely required during a specific developmental period for courtship, and plays a minor role during adulthood; secondly, the sex circuitry without *fru^M^* or with different levels of *fru^M^* has different properties such that males would have experience-dependent courtship acquisition, or innate courtship but with different sexual orientation. Such flexibility of the sex circuitry is tuned by different *fru^M^* expression. Red circles and blue triangles represent corresponding *fru^M^* levels and courtship levels (red: male-male courtship; blue: male-female courtship).

## Discussion

Previous findings show that *fru^M^* expression commences at the wandering third instar larval stage, peaks at the pupal stage, and thereafter declines but does not disappear after eclosion (Lee et al., 2000), which suggests that *fru^M^* may function mainly during development for adult courtship behavior despite of no direct evidence. Here we temporally knocked down *fru^M^* expression in different developmental stages for 2 days and found that males with *fru^M^* knocked down during pupation rarely courted, while males with *fru^M^* knocked down during adulthood courted normally towards females. This is the first direct evidence that *fru^M^* is required during development but not adulthood for courtship behavior. However, we also found a minor role of *fru^M^* during adulthood in suppressing male-male courtship, as males with *fru^M^* knocked down or *tra* overexpressed during adulthood displayed slightly enhanced male-male courtship or male chaining behaviors. Note that a previous study found that removal of *transformer 2 (tra2)* specifically during adulthood using a temperature sensitive *tra2* allele induced 8 out of 96 females to show male-type courtship behaviors (Belote and Baker, 1987), which suggests that expression of FRU^M^ and DSX^M^ (by removal of *tra2* function in females) during adulthood is sufficient to masculinize CNS to some extent and induce a small fraction of females to display courtship behaviors. A recent study also found that *fru^M^* expression in the *Or47b*-expressing olfactory receptor neurons depends on the activity of these neurons during adulthood (Hueston et al., 2016). Based on all these findings, we propose that *fru^M^* expression during pupation is crucial for specifying a sex circuitry that allows innate courtship towards females, and its expression during adulthood may be activity-dependent in at least some neurons and modulates some aspects of courtship *(e.g.,* inhibits male-male courtship). Thus, there are at least two separate mechanisms that *fru^M^* contributes to the sex circuitry, one during a critical developmental period to build the female-directed innate courtship into that circuity, and the other during adulthood to modulate neuronal physiology in an experience-dependent manner.

Most importantly, we revealed striking flexibility of the fly sex circuitry by manipulating *fru^M^* expression. We listed four cases with *fru^M^* manipulation here for comparison: (1) males with a sex circuitry having wild-type *fru^M^* function have innate heterosexual courtship, as they court readily towards females, but do not court males no matter how long they meet; (2) males with a sex circuitry having no *fru^M^* function lose the innate courtship ability, but have the potential to acquire courtship towards males, females, and even other species in an experience-dependent manner; (3) males with a sex circuitry having limited *fru^M^* expression *(e.g.,* 40%) have innate homosexual courtship, as they court readily towards other males, but rarely court females; (4) males with a sex circuitry having limited *fru^M^* expression outside brain (but intact *fru^M^* expression in brain) are innately bisexual, as they court equally towards females or males. Although previous studies found that different *fru^M^* alleles (*e.g.*, deletions, inversions or insertions related to *fru*) showed very different courtship abnormalities (Anand et al., 2001; Villella et al., 1997), it was very hard to link *fru^M^* function to the flexibility of sex circuitry, and often seen as allele-specific or background-dependent phenotypes. Our study using relatively simple genetic manipulations that generate dramatical different courtship behaviors promoted us to speculate a different view about the role of *fru^M^*. instead of simply being a master gene that controls all aspects of male courtship, *fru^M^* is not absolutely necessary for courtship, but changes the wiring of the sex circuitry during development such that the sex circuitry may function in very different ways, ranging from innately heterosexual, homosexual, bisexual to largely experience-dependent acquisition of the behavior. Such flexibility of the sex circuitry is tuned by different *fru^M^* expression, such that changes of *fru^M^* regulatory regions during evolution would easily select a suitable functional mode of the sex circuitry.

## Materials and methods

### Fly Stocks

Flies were maintained at 22 or 25°C in a 12 h: 12 h light:dark cycle. Canton-S flies were used as the wild-type strain. Other stocks used in this study include the following: *fru^GAL4^* (Stockinger et al., 2005), *UAS-fruMi* and *UAS-fruMiScr* (Meissner et al., 2016), *fru^LexA^* and *fru^4-40^* (Pan and Baker, 2014) and *Otd-Flp* (Asahina et al., 2014). *UAS-traF* (BL#4590), *tub-GAL80^ts^* (BL#7019), *tub>GAL80>* (BL#38881) and *tub>stop>GAL80* (BL#39213) were from Bloomington Drosophila Stock Center.

### Courtship and Chaining Assays

For the single-pair courtship assay, the tester males and target flies (4-8 days old) were gently aspirated into round 2-layer chambers (diameter: 1 cm; height: 3 mm per layer) and were separated by a plastic transparent barrier that was removed ~30 min later to allow courtship test. Courtship index (CI), which is the percentage of observation time a fly performs any courtship step, was used to measure courtship to female targets or between 2 males. Paired male-male courtship used 2 males of the same genotype but focused on the male fly that first initiated courtship (courtship of the initiator to the other). All tester flies were single housed if not otherwise mentioned. Each test was performed for 10 min.

For male chaining assay, tester males (4-8 days old) were loaded into large round chambers (diameter: 4 cm; height: 3 mm) by cold anesthesia. Tests were performed daily for 4 consecutive days (3 hours after grouping as day 0, then days 1–3). Chaining index (ChI), which is the percentage of observation time at least 3 flies engaged in courtship together, was used to measure courtship in groups of 8 males.

### Immunohistochemistry

We dissected brains and ventral nerve cords of 5-7 days old males in Schneider’s insect medium (Thermo Fisher Scientific, Waltham, MA) and fixed in 4% paraformaldehyde (PFA) in phosphate-buffered saline (PBS) for 30 min at room temperature. After washing four times in PAT (0.5% Triton X-100 and 0.5% bovine serum albumin in PBS), tissues were blocked in 3% normal goat serum (NGS) for 60 min, then incubated in primary antibodies diluted in 3% NGS for ~24 hr at 4°C, washed (4 × 15-min) in PAT at room temperature, and incubated in secondary antibodies diluted in 3% NGS for ~24 hr at 4°C. Tissues were then washed (4 × 15-min) in PAT and mounted in Vectorshield (Vector Laboratories, Burlingame, CA) for imaging. Primary antibodies used were rabbit anti-GFP (1:1000; A11122, Invitrogen, Waltham, MA) and mouse anti-Bruchpilot (1:50; nc82, Developmental Studies Hybridoma Bank, Iowa City, IA). Secondary antibodies used were goat anti-mouse IgG conjugated to Alexa 555 (1:500, A28180, Invitrogen) and goat anti-rabbit IgG conjugated to Alexa 488 (1:500, A11008, Invitrogen). Samples were imaged at 10× or 20× magnification on a Zeiss 700 confocal microscope and processed with ImageJ.

### Statistics

Experimental flies and genetic controls were tested at the same condition, and data are collected from at least two independent experiments. Statistical analysis is performed using GraphPad Prism and indicated inside each figure legend. Data presented in this study were first verified for normal distribution by D’Agostino-Pearson normality test. If normally distributed, Student’s *t* test is used for pairwise comparisons, and one-way ANOVA is used for comparisons among multiple groups, followed by Tukey’s multiple comparisons. If not normally distributed, Mann-Whitney U test is used for pairwise comparisons, and Kruskal-Wallis test is used for comparisons among multiple groups, followed by Dunn’s multiple comparisons.

## Acknowledgements

We thank the Bloomington Drosophila Stock Center for fly stocks. This work was supported by grants from National Key R&D Program of China (2019YFA0802400), the National Natural Science Foundation of China (31970943 and 31622028), and the Jiangsu Innovation and Entrepreneurship Team Program.

---

**Figure 3—figure supplement 1.**
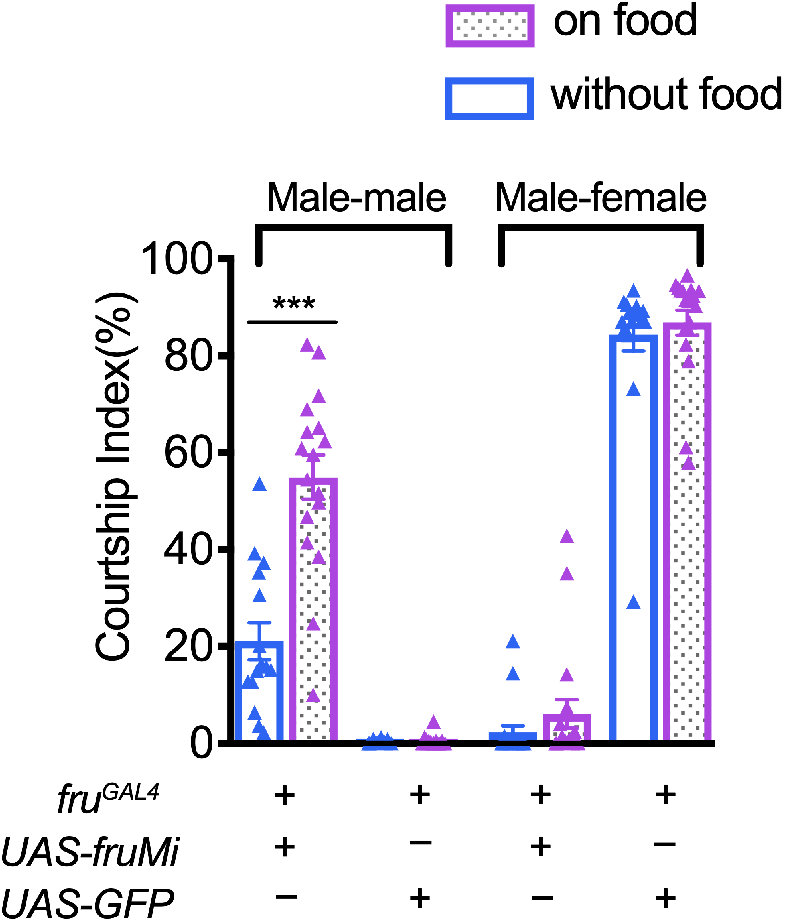
Comparison of male courtship with or without food. Male-male courtship in *fru^M^* knocked down males is higher in the presence of food. *n* = 15, 17, 15, 18, 18, 18, 18 and 18 from left to right respectively. \*\*\**p* < 0.001, unpaired t-test. Error bars indicate SEM.

